# Simulating Rumen Conditions using an Anaerobic Dynamic Membrane Bioreactor to Enhance Hydrolysis of Lignocellulosic Biomass

**DOI:** 10.1101/2023.02.20.529314

**Authors:** Xavier Fonoll, Kuang Zhu, Lucy Aley, Shilva Shrestha, Lutgarde Raskin

## Abstract

An anaerobic dynamic membrane bioreactor (AnDMBR) was used to mimic rumen conditions to enhance the hydrolysis of lignocellulosic materials and the production of volatile fatty acids (VFA) when treating food waste. The AnDMBR was inoculated with cow rumen content and operated at a 0.5-day hydraulic retention time, 2 to 4-day solids retention time, a temperature of 39 °C, and a pH of 6.3, characteristics similar to those of a rumen. An operational strategy of repeating settle-permeate-mix cycles was utilized to simulate the regurgitation of ruminant animals. Removal rates of neutral detergent fiber (NDF) and acid detergent fiber (ADF) of 58.9 ± 8.4 % and 69.0 ± 8.6 %, respectively, and a VFA yield of 0.55 ± 0.12 g VFA/g COD_fed_ were observed when operating at an organic loading rate of 18 ± 2 kg VS/m^3^/day. Sequencing results of both 16S rRNA gene and 16S rRNA revealed that a microbial community resistant to both disruption and reinoculation developed six weeks after inoculation. Up to 66.7 ± 5.7 % of the active microbial populations (based on 16S rRNA sequence data) and 51.0 ± 7.0 % of the total microbial populations (based on 16S rRNA gene sequence data) present in the rumen-mimicking AnDMBR originated from the rumen inoculum. This study offers a reliable strategy to leverage the unique features of a rumen; the rumen-mimicking AnDMBR achieved high hydrolysis and acidogenesis rates even when treating substrates drastically different from those fed to ruminants.

## INTRODUCTION

Organic waste generated in domestic, industrial, and agricultural activities is a rich reservoir for resources, including energy and materials, that can be recovered to replace their fossil-sourced counterparts. One of the key technologies for resource recovery, anaerobic digestion (AD), has been used to treat wastes and recover resources from wastes around the world for centuries and it started to be widely implemented for the treatment of diverse waste streams around 30 years ago.^1,2^ However, the implementation of AD remains challenging for a range of applications given the high capital and operating costs and relatively low product yields for conventional wet digesters.^3,4^ One of the main drawbacks of AD is the slow hydrolysis of recalcitrant lignocellulosic materials, requiring high retention times, large reactor footprints, and low methane yields^3,5,8,79^. Lignocellulosic materials are comprised of cross-linked structures of cellulose, hemicellulose and lignin that are difficult to break down by anaerobic microorganisms and are present in high amounts in various wastes (agricultural wastes, yard and park wastes, vegetable and fruit fraction of food wastes, paper waste, etc.)^7,8,10^. Combining substrate pretreatment (chemical, mechanical, or enzymatic) with long solids retention times (SRT>40 days) can hydrolyze up to 90% of lignocellulose. Novel bioreactor designs, such as anaerobic dynamic membrane bioreactors (AnDMBR), can reduce footprint requirements by decoupling the SRT and the hydraulic retention time (HRT). Nevertheless, operating costs remain high due to energy and chemical requirements^11–16^.

Microbial communities in natural environments like the rumen, the stomach of ruminant animals, are capable of effectively hydrolyzing lignocellulosic materials^10,17^. In fact, the bacterial and fungal communities in the rumen can hydrolyze up to 62% of the lignocellulosic components without pretreatment.^18–23^. The rumen has an organic loading rate (OLR) between 65 and 140 g TS L_R_^−1^ day^−1^, an SRT of 30 to 90 h, and an HRT of 5 to 20 h.^24–28^ The SRT is longer than the HRT because there is a narrow passage between the first stomach (rumen) and second stomach (reticulum) that promotes the formation of a biofilm where food collects while trying to pass from the rumen to the reticulum.^26,28^ This retention of solids allows lignocellulose to be retained in the rumen long enough to be hydrolyzed.^24^ Once lignocellulosic components are hydrolyzed and the particle size reduced, solubilized compounds and smaller particles are fermented to produce volatile fatty acids (VFAs). VFAs are removed through the walls of the rumen to provide energy to the host and the rest of the hydrolysis and fermentation products travel to the reticulum.^26,29^ Previous studies have utilized rumen content as the inoculum or a co-substrate and recreated rumen hydraulic conditions to enhance hydrolysis rates of lignocellulosic substrates^23–38^. Nevertheless, it is unclear if this strategy is feasible to treat other common organic waste streams substantially different from the feed of ruminants (e.g., food waste), because the substrate characteristics are a crucial shaping factor for the structure and function of microbial communities.^10,31^ Another potential shaping factor of rumen function that was not previously considered is the regurgitation and mastication by ruminant animals that can break biofilms to avoid inhibiting the growth of rumen hydrolytic microorganisms^46^. The formation of biofilms in the rumen is crucial to enable the fast hydrolysis of lignocellulose; however, it may also create unfavorable conditions for hydrolytic microbes. Biofilms developed in the rumen can trap hydrolytic and acidogenic microorganisms in an unfavorable local environment with high concentrations of inhibitors like VFAs and low pH.^46^ Mason & Stuckey (2016) hypothesized that one of the functions of regurgitation is to avoid microbial toxicity in the biofilm. During regurgitation, biofilm particles travel from the rumen into the mouth and are chewed and broken down by the ruminant’s teeth. Moreover, the pH in the saliva is 8.0-8.8, which helps bring the pH levels in these particles to a more desirable level.^47^ After regurgitation, the remaining lignocellulosic components and microorganisms are sent back into the rumen where the formation of a biofilm and hydrolysis restart.

An AnDMBR was designed and operated to mimic rumen conditions by inoculating the system with cow rumen content, selecting the HRT, SRT, temperature, and pH similar to those of a cow rumen, and simulating regurgitation behavior by disrupting the dynamic membrane (biofilm). The goal of this study was to evaluate the ability of this system to retain the rumen microbial community, enhance the hydrolysis rate of lignocellulosic materials, and maintain microbial and performance resiliency during operational disruptions when treating food waste.

## MATERIALS AND METHODS

### Inoculum and substrate

The inoculum consisted of a mixture of liquid and solid rumen content collected separately from a fistulated cow at a Michigan State University dairy farm (Lansing, MI). The rumen samples were kept at 39°C using a warming container (Wagan EL6224) during the 1-h transportation time and were used immediately upon arrival in the laboratory for inoculation. The inoculum was prepared by mixing 6.2 L of liquid rumen content with 300 g of rumen solids. The inoculum used to seed the bioreactor was characterized immediately after mixing the liquid and solid rumen content using methods described below.

Post-consumer food waste collected from a dining hall at the University of Michigan (Ann Arbor, MI) was used as the substrate. It contained unconsumed grains, fruits, vegetables, meat, and sauce mixed with napkins and tea bags and was processed with a pulper that grinded and dewatered the food waste. The food waste slurry was stored at −20 °C. When ready to be used, it was thawed overnight at 4 °C and blended (Waring Commercial MX1200XTX 3.5 HP) with deionized water to reach a total solids (TS) concentration of approximately 75 g TS L^−1^ to ensure adequate mixing during bioreactor operation. This final food waste mixture was characterized using methods described below.

### Rumen-mimicking AnDMBR and operating conditions

A 6.2 L (working volume) semi-continuous AnDMBR controlled remotely by LabVIEW (National Instruments, Austin, TX, USA) data acquisition software was utilized to recreate rumen conditions. The rumen-mimicking AnDMBR is referred to as rumen AnDMBR throughout this study. A dynamic membrane, a cake layer comprised of biofilm and solids, was established on a 100 μm pore size mesh support (area of 314 cm^2^) to achieve solid-liquid separation, allowing the recreation of rumen hydraulic conditions (Figure S1). Deionized water was continuously fed into the bioreactor after the daily manual feed of food waste to ensure an HRT of 12 h. Suspended biomass was removed from the bioreactor bulk solution every other day before feeding to control the SRT. The total suspended solids (TSS)-based SRT was 97 ± 39 h while the total solids (TS)-based SRT was 51 ± 15 h.

The pH in the rumen AnDMBR was continuously monitored by a pH electrode (SL 83 SI analytics) and maintained between 6.2-6.3 by automatically dosing 1-M NaOH solution through a peristaltic pump (Masterflex L/S). A cyclic operational strategy was implemented to simulate rumen conditions (Figure S1): i) the mixing stops for 49 min during the settling stage to generate a dynamic membrane on the stainless-steel mesh, similar to the biofilm formation taking place between the rumen and the reticulum; ii) the permeate was extracted through the dynamic membrane for 1 min, simulating the passage of hydrolysis products from the rumen into the reticulum; and iii) the bioreactor was mixed at 100 rpm to allow NaOH dosage for pH control (NaOH can only be dosed during the mixing stage to ensure accurate pH reading from the pH probe) and to simulate the regurgitation and mastication. Because the highest VFA production rate was expected to occur immediately after the daily feeding, the bioreactor was continuously mixed for 3 hours at 100 rpm after feeding to avoid over-acidification during the settling and permeating stages. The temperature in the bioreactor was controlled at 39 °C, similar to a rumen, using a recirculating water bath (Polyscience, Niles, IL, USA). A 5 L Tedlar gasbag was connected to the headspace of the bioreactor for biogas production measurements. During the 110-day operation, the OLR was maintained at 18 ± 2 g volatile solids (VS) L_R_^−1^ day^−1^. On Day 44, an over-dosing of NaOH caused the pH in the bioreactor to increase above 12 for 4 hours. With the assumption that the incident with NaOH on day 44 could have affected the activity of rumen microbial populations, the bioreactor was reinoculated with rumen content on Day 79 to bring back these populations into the system.

### Chemical analyses

The inoculum was characterized at the time of inoculation (day 0 and day 79) and substrate samples were analyzed once a week. Suspended biomass and permeate samples were collected three times a week. TS, VS, TSS, and volatile suspended solids (VSS) were analyzed following standard methods.^48^ Total, partial, and intermediate alkalinity were determined using a titration method as described by Ripley et al.^49^ The neutral detergent fiber (NDF) and acid detergent fiber (ADF) analyses were carried out in an ANKOM220 Fiber Analyzer (ANKOM Technology, Macedon, NY, USA) following manufacturer’s protocols.^50^ Concentrations of VFAs from C2 to C8, including iso-forms of C4 and C5, were determined using an Agilent Technologies 7890B gas chromatograph (Santa Clara, CA) equipped with a Stabilwax-DA column (Restex) and a flame ionization detector. The oven temperature was held at 55 °C for 1 min, then increased to 205 °C at 10 °C min^−1^ and held at 205 °C for 8 min. Injector and detector temperatures were set to 250 °C and 300 °C, respectively, and nitrogen was used as the carrier gas. The total and soluble chemical oxygen demand (tCOD and sCOD) analyses were performed using Lovibond™ medium-range (0– 1500 mg L^−1^) COD digestion vials (Tintometer, Germany). Before performing VFA and sCOD analyses, all samples were centrifuged at 10,000 g for 10 min and the supernatant was filtered through 0.45 μm nylon membrane filters (TISCH Scientific, North Bend, OH, USA). Phosphoric acid was used to adjust the sample pH below 2 before the VFA analysis.

The gas produced in the rumen AnDMBR was collected in Tedlar bags and measured daily by using a 100 mL gas-tight glass syringe. The gas composition (CH_4_, H_2_ and CO_2_) was measured three times a week using a gas chromatograph (Gow-Mac, Bethlehem, PA) coupled with a thermal conductivity detector (GC-TCD). The temperature of the column, injector, and detector were set to 104 °C, 80 °C, and 115 °C for CH_4_, H_2_ and CO_2_, respectively, and the current was set at 120 mA. Hydrogen was used as the carrier gas for carbon dioxide and methane measurements while nitrogen was used as the carrier gas to measure hydrogen. Standard gas samples (ShopCross, Greensboro NC, USA) with varying mixtures of methane, carbon dioxide, and hydrogen were used for GC-TCD calibration. Suspended biomass, permeate, and substrate samples collected on days 90, 95, 100 and 107 were shipped to Eurofins (Horsham, PA) for protein and lipid analyses. Protein analysis was performed using AOAC 990.03 and AOAC 992.15 combustion methods and lipid analysis was performed using AOAC 948.22 Sohxlet extraction method.

### Biomass sampling, DNA extraction, and 16S rRNA and 16S rRNA gene sequencing

Biomass samples were pelletized by centrifugation at 11,000 g for 10 min at 4°C immediately after collection (suspended biomass and permeate samples) or preparation (food waste sample) and inoculation (inoculum sample). After centrifugation, the supernatant was decanted, and the pellet was immediately stored at −80°C until DNA and RNA extraction.

DNA was extracted using a cetyl trimethylammonium bromide (CTAB) method^51^ with an additional bead-beating step (Mini-Beadbeater-96, BioSpec Products, Bartlesville, OK, USA) for 1.5 min using 0.1 mm diameter zirconium beads. DNA quality and quantity were determined by spectrophotometry (Nanodrop 1000, Thermo Fisher Scientific, Wilmington, DE, USA) and Qubit 2.0 Fluorometer (Invitrogen, Life Technologies, MA, USA).

RNA was extracted with the TRIzol method using the Zymo Direct-zol RNA kit (Zymo). DNA/RNA shield reagent (Zymo) was added to the cell pellets at a 1:10 pellet to reagent ratio (w:w) immediately prior to bead beating while the samples were still frozen to prevent RNA degradation. Baked (450 °C) zirconium beads with diameters of 0.5 m and 0.1 mm were mixed at a 7:6 ratio (w:w) and used for bead beating. After bead beating for 1.5 min, the RNA was extracted following the manufacturer’s protocol without DNA digestion. RNA quality was determined by a bioanalyzer (Biorad, RNA Stdsense kit, Hercules, CA, USA). The RNA was reverse transcribed to cDNA with a SuperScript IV VILO kit (Invitrogen, MA, USA) following the manufacturer’s protocol with DNA digestion. The quantity of cDNA was determined with Qubit 2.0 Fluorometer (Invitrogen, Life Technologies, MA, USA).

DNA and cDNA samples were submitted to the Host Microbiome Initiative (University of Michigan, Ann Arbor, MI, USA) for 16S rRNA gene and 16S rRNA sequencing via Illumina MiSeq (San Diego, CA, USA). Food waste samples were not sequenced because no RNA could be extracted as indicated by electrophoresis results (Figure S13). Universal primers F515 and R806 targeting the V4 region of the 16S rRNA gene were used for PCR amplification.^52^ The sequences obtained were processed using mothur (version 1.45.3) following the MiSeq SOP.^53^ The SILVA database (Release 138.1) was used for taxonomic classification and Amplicon Sequence Variant (ASVs) that could not be classified at the genus level were denoted as “unclassified” followed by the name of the lowest taxonomic group in which the ASV could be classified. The closest relatives of the abundant (>1%) ASVs were determined using a BLAST analysis.

For the quantification of *Neocallimastigomycota* in the inoculum and suspended biomass samples, a qPCR analysis was performed following the protocol described by Dollhofer et al. (2016).^54^

### Statistical analyses

Beta diversity was represented in a two-dimensional graph using nonmetric multidimensional scaling (NMDS) analyses based on Bray-Curtis dissimilarity index as implemented in mothur. The statistical differences in the composition of the microbial community (total microbial populations determined by 16S rRNA gene sequencing and active microbial populations determined by 16S rRNA sequencing) among different samples was tested with an analysis of molecular variance.^53^

## RESULTS AND DISCUSSION

### The rumen AnDMBR produced a high VFA yield at a high OLR

The VFA yield in the rumen AnDMBR stabilized in the range of 0.55 – 0.56 g VFA as COD g VS_fed_^−1^ shortly after inoculation (Figure 1, Table S2). The concentrations of acetate, propionate, and butyrate in the permeate comprised 35%, 32%, and 18% of the total VFAs (Table S2).

**Figure 1.**
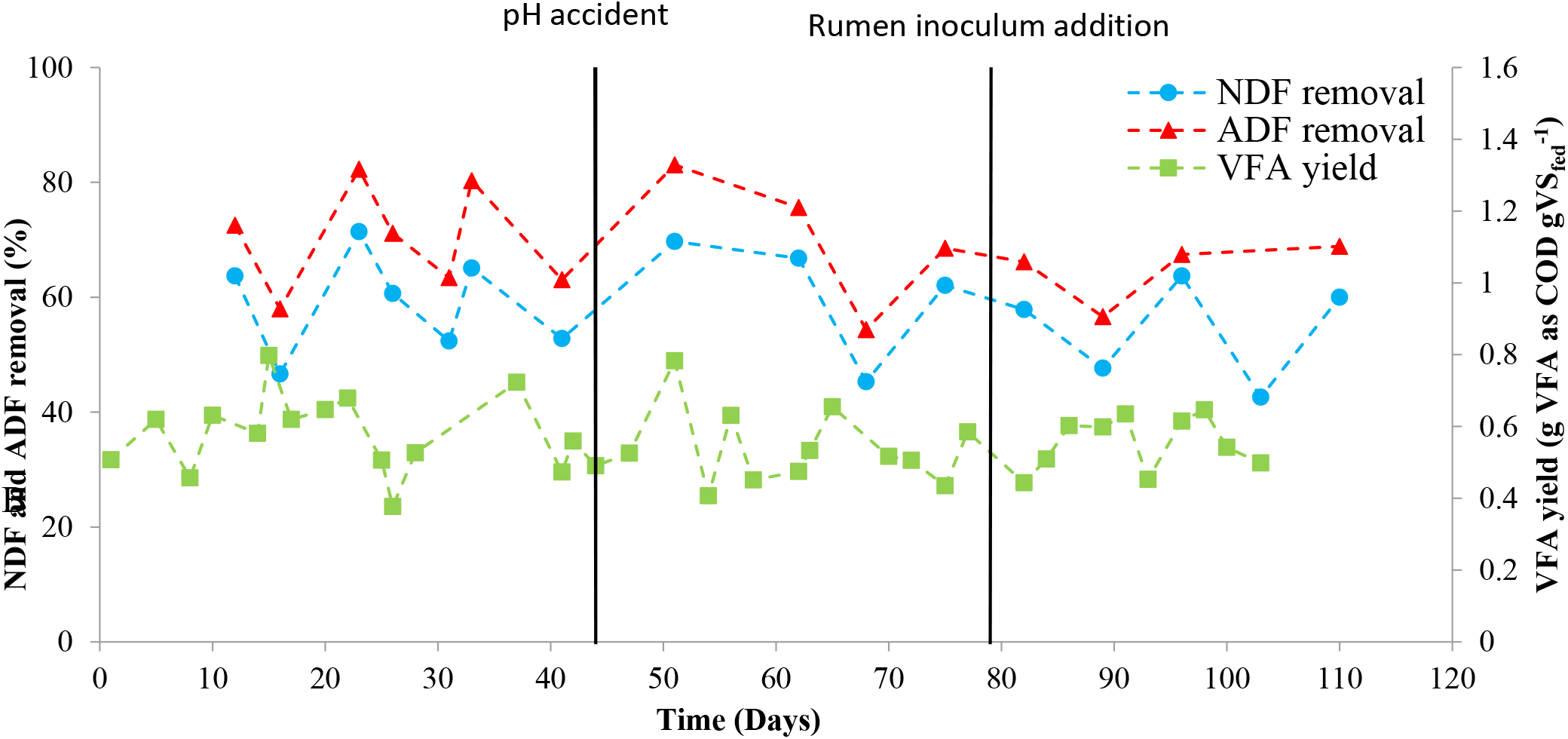
NDF and ADF removal and VFA yield in the rumen AnDMBR. The vertical lines indicate the increase of pH for 4 hours (day 44) and the addition of rumen inoculum (day 79).

The VFA yields in the rumen AnDMBR were comparable with other fermentation systems treating food waste but the rumen AnDMBR had a much lower HRT of 12 ± 2 h and a higher OLR of 18 ± 2 g VS L_R_^−1^ day^−1^ (Table 1), supporting our goal of designing a system with a small reactor footprint and high treatment capacity.

**Table 1.** Performance comparison between the rumen AnDMBR developed in this study and other rumen-simulating bioreactors

|  | This study | Other studies.<sup>55–61</sup> |
| --- | --- | --- |
| VFA yield (g VFA g VS<sub>fed</sub><sup>-1</sup>) | $0.38 \pm 0.10$ | 0.3 – 0.5 |
| HRT (days) | $0.50 \pm 0.08 \text{ h}$ | >5 |
| OLR (g VS L<sub>R</sub><sup>-1</sup> day<sup>-1</sup>) | $18 \pm 2$ | <11 |

For conventional continuous stirred-tank reactors, high OLRs are typically achieved by high feed concentrations, resulting in high VFA concentrations that can inhibit hydrolytic and acidogenic microbial populations and decrease VFA yields^55,56,26,62,63^. The rumen AnDMBR, however, achieved a high OLR by operating at a very low HRT and permeate VFA concentrations were low (6.0 ± 1.9 g VFA as COD L^−1^, Table S2 and Figure S2B), allowing for high VFA yields.^26^

The high VFA yields for the rumen AnDMBR may also be explained by the efficient hydrolysis of lignocellulosic materials. The removal rate for NDF, which represents hemicellulose, cellulose, and lignin, was 58.9 ± 8.4 % and the removal rate for ADF, which represents cellulose and lignin, was 69.0 ± 8.6 %. These NDF removal rate was at the high end of the ranges measured in real rumens (25-62% of NDF removed).^18–23^ Nonetheless, these encouraging results do not necessarily translate to hydrolysis kinetics comparable to those in real rumens because of the drastic difference in substrate characteristics (Table S1). Therefore, given the high concentrations of lipids and proteins in food waste, their removal rates were also analyzed. A high lipid removal rate of 90.0 ± 3.8 % was observed, while 42.0 ± 3.8% of proteins were solubilized (Table S2). Overall, the rumen AnDMBR presented one of the highest lignocellulose removal rates and VFA yields, and the results are similar to those observed by Gijzen et al.^39^, who reported a rumen AnDMBR with an HRT and SRT close to those reported for real rumens but did not replicate the regurgitation effect (Table S3). In the rumen AnDMBR, the cyclic settle-permeate-mix operation consistently achieved SRT and HRT decoupling while maintaining high permeation fluxes through the dynamic membrane (415 ± 43 LMH). This was because at each cycle, a new dynamic membrane resulting in high permeability was established during the settle stage after the previous one was disrupted during the mixing stage. The dual-purpose mixing and dynamic membrane disruption only took place for 10 min every hour, requiring less energy input than continuous mixing. An even shorter mixing stage can be explored in future work to further reduce energy requirements.

An accidental over-dose of NaOH occurred on day 44, which increased the pH to 12 for four hours. The VFA yields stayed consistently high while the ADF removal decreased from 83.0 % to 54.4 % and the NDF removal decreased from 69.7 to 45.3 % after the NaOH overdose (Figure 1). The continuous decrease in NDF and ADF removal from day 50 to day 68 brought the assumption that the NaOH incident affected rumen microbial populations involved in hydrolysis. On day 79, half of the content of the bioreactor was replaced by a fresh rumen inoculum in an attempt to recover the rumen microbes involved in hydrolysis. However, the NDF and ADF removals had already increased to 62.0 % and 68.5 % on day 75, before re-inoculation.

### The rumen AnDMBR supported a resilient and diverse microbial community

The relative activities (as determined by 16S rRNA sequencing) of the dominant microbial populations in the rumen AnDMBR throughout the 110-day operating period are shown in Figure 2. Similar data derived from 16S rRNA gene sequencing are presented in Figure S3.

**Figure 2.**
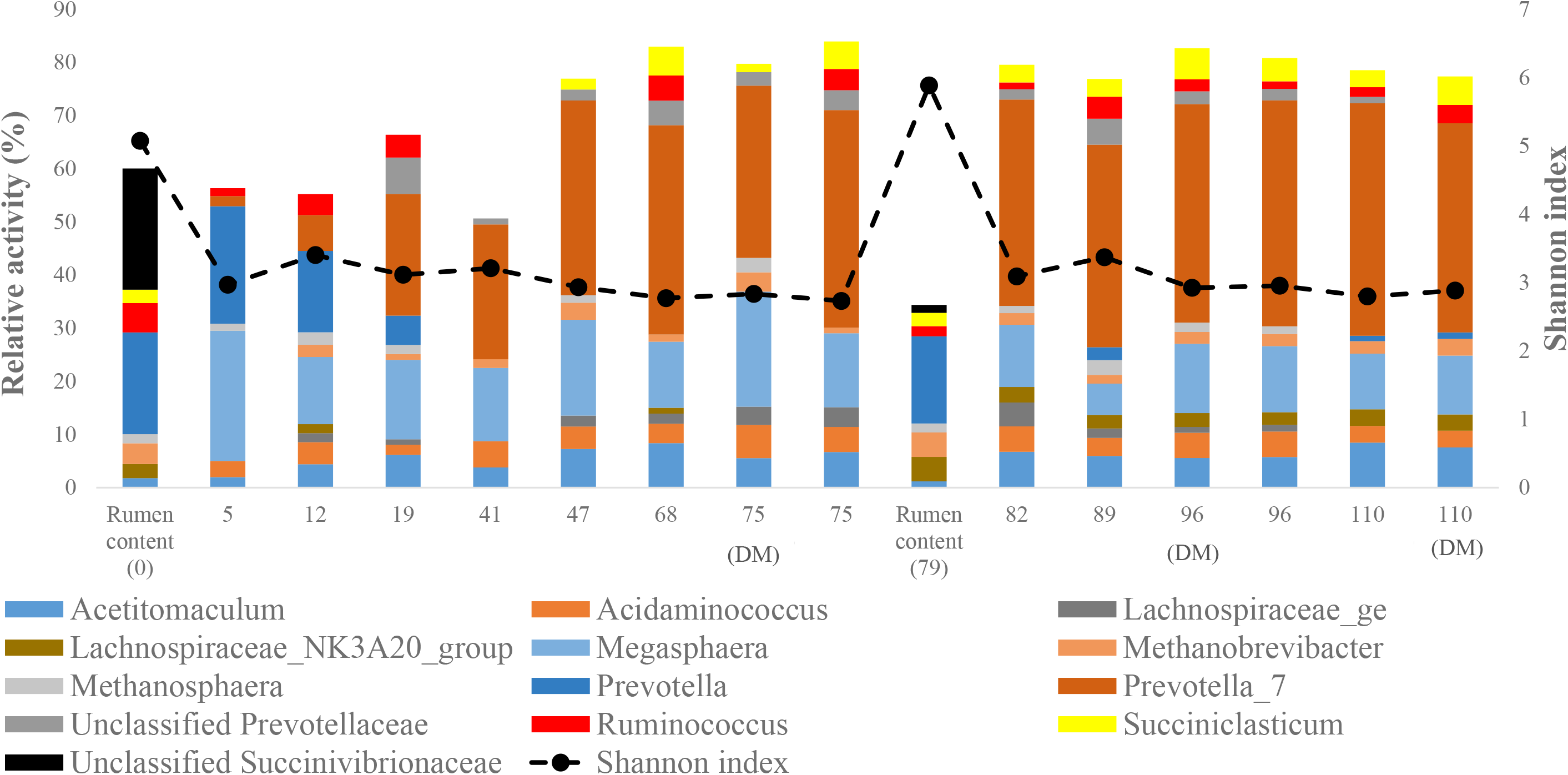
Relative activities (determined by 16S rRNA sequencing) of dominant genera in the inoculum and bioreactor suspended biomass over time. Only genera present at relative activities greater than 1% in at least 60% of the samples (n=10) or present at a relative activity of 15% at least once are shown. The Shannon diversity index was calculated considering all the ASVs. DM indicates that the samples were taken from the dynamic membrane.

*Prevotella 7*, *Megasphaera*, *Acidaminococcus*, and *Acetitomaculum* were the most active and abundant populations (Figure 2 and Figure S3). *Prevotella 7* belongs to the family *Prevotellaceae* species of which are found in high abundance in the rumen and can hydrolyze plant biomass or perform fermentation of sugars into VFAs.^64–67^ *Megasphaera* is a genus involved in the production of VFAs from lactate, glucose, or succinate.^68^ *Acidaminococcus* uses amino acids from the hydrolysis of proteins to produce VFAs^69,70^. *Acetitomaculum* is a genus involved in reductive acetogenesis and uses H_2_ and CO_2_ to produce acetate, thus competing with hydrogenotrophic methanogens for substrates.^71^ The high activity of *Acetitomaculum* (2.0–8.5 %) may explain the low methane yield in the rumen AnDMBR (2.5 ± 1.2 g CH_4_ g COD_fed_^−1^). The accidental over-dose of NaOH on day 44, which increased the pH to 12 for four hours, and the subsequent re-inoculation on day 79 did not have an apparent impact on the overall microbial community structure in the rumen AnDMBR (Figure 2 and Figure S3), indicating a highly resilient microbial community.

A goal for this research was for the rumen AnDMBR to retain key hydrolytic rumen bacterial genera like *Fibrobacter* and *Ruminococcus*, and hydrolytic fungi like *Neocallimastigomycota*.^65,72^ Overall, the genus *Ruminococcus* stayed highly active with a relative activity of 1.3 - 4.7% (Figure 3). A closer investigation revealed that most of the ASVs comprising the *Ruminococcus* genus belonged to *Ruminococcus flavefaciens* and *Ruminococcus albus*, cellulolytic and hemicellulolytic species (Figure S4 and Table S4).^65,67^ *Fibrobacter*, an important fibrolytic genus in the rumen,^31,73^ reemerged after Day 75 and maintained its relative activity and abundance after reinoculation (Figures S5). Despite the fluctuating relative activity of *Fibrobacter*, high and consistent hydrolysis rates were observed throughout the experiment, suggesting that it is not a rate-dominating factor in hydrolyzing food waste. The reason for this might be that food waste is less lignocellulosic than ruminants’ food (e.g., alfalfa). *Neocallimastigomycota*, an important hydrolytic anaerobic fungus in cow rumen, was washed out (Figure S6), potentially due to the high readily biodegradable carbohydrate concentration repressing expression of genes encoding for hydrolytic enzymes and the disruption of the dynamic membrane on which the fungi rhizoid can attach.^74,75^ Future work is needed to validate these hypotheses.

**Figure 3.**
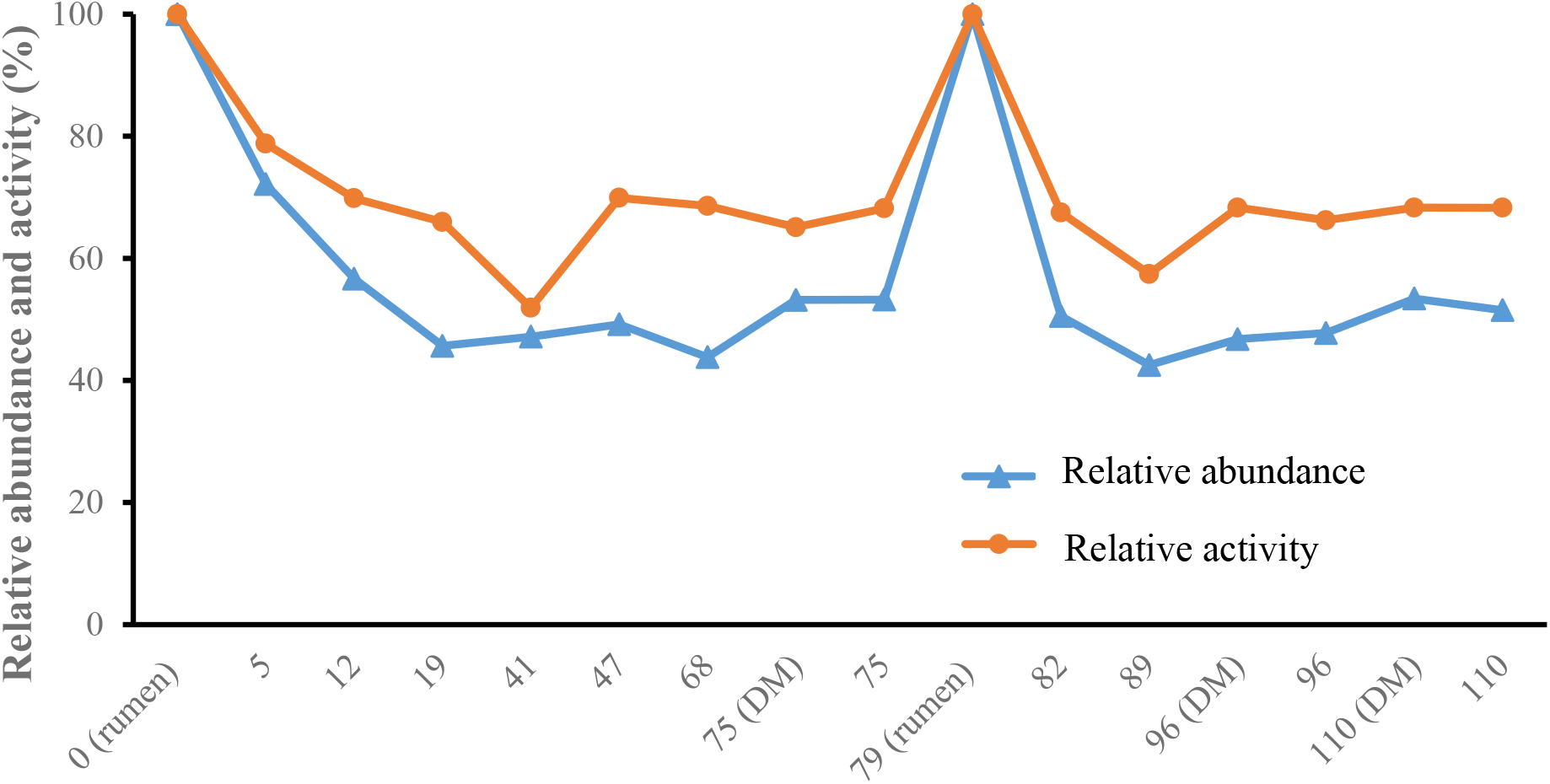
Proportion of ASVs present in rumen inocula that were maintained in the rumen AnDMBR over time. The ASV from the rumen content used on day 0 were considered for the samples between days 5 to 75 and the ASV from the rumen content used on day 79 were considered for the samples between days 82 to 110.

DNA and RNA were also extracted from permeate samples and the most abundant and active microorganisms in the permeate were *Acetitomaculum*, *Acidaminococcus*, *Lachnospiraceae NK3A20*, *Megasphaera*, Methanobrevibacter, *Prevotella_7*, unclassified *Prevotellaceae*, *Succiniclasticum* and *Ruminococcus* (Figure S7 and S8). The microbial community structures (based on relative abundance and activity) in the digestate, dynamic membrane and permeate were not statistically significantly different (p=0.402 for relative activity and p=0.185 for relative abundance) (Figure S9).

### The rumen AnDMBR permeate provides versatile downstream treatment options

Only 2.5% of the influent COD was converted into methane in the rumen AnDMBR (Table S2 and Figure S2A). With a highly hydrolyzed stream containing a variety of readily available substrates in the permeate, the rumen AnDMBR is an ideal first phase bioreactor for a variety of downstream processes to recover valuable resources from waste.^76–78^ A common strategy for two-phase treatment involves biogas production using a methanogenic second-phase anaerobic digester^79,80^. The microbial community present and active in the permeate with diverse functions can offer resource recovery options other than biogas. For example, *Megasphaera*, a genus with high relative activity, was mostly represented in the rumen AnDMBR by *Megasphaera elsdenii* (Figure S10 and Table S5) which is a rumen chain-elongation species involved in the production of butyrate and caproate from lactate^68,81^. In fact, the permeate from this study was successfully used as a substrate for chain elongation in a continuous system in which *Megasphaera* was enriched.^76^

### The rumen environment was sufficiently replicated to efficiently hydrolyze food waste

Figure 3 shows the proportion of populations in the microbial community in the rumen AnDMBR that originated from the rumen inocula.

An average 51.0 ± 7.0 % of the ASVs of the microbial community in the rumen AnDMBR were present in the rumen inoculum. This proportion was higher (66.7 ± 5.7 %) for the active microbial populations. In contrast, when Fonoll et al. (2021) used rumen content as a co-substrate, only 36% of the ASVs present in the 39°C semi-continuous, neutral-pH bioreactor were also present in the rumen and no enhancement of hydrolysis was observed^31^. This suggests the importance of recreating rumen conditions, including HRT, SRT, and pH, beside introducing rumen microbiome when the goal is to use rumen microorganisms to enhance hydrolysis. However, the microbial community in the rumen AnDMBR were less diverse compared to the rumen inoculum as indicated by the decreased Shannon Diversity Indices, consistent with other studies simulating cow rumen (Figure 2 and S3).^40,42,76^ Moreover, the microbial community structure and activity in the rumen AnDMBR were statistically significantly different from the one in the rumen content (p=0.003 for relative activity and p=0.014 for relative abundance) (Figure 4 and S11). For example, *Prevotella* and *Succinivibrionaceae*, who were abundant and active in the rumen inoculum gradually disappeared from the rumen AnDMBR (Figure 2 and S3). On the other hand, *Prevotella 7*, *Megasphaera*, *Acidaminococcus*, and *Acetitomaculum*, who had low abundances and activities in the rumen inoculum became the most active and abundant genera (Figure 2 and S3). The shifts are likely caused by the difference in substrate characteristics. Ruminant feed usually has high lignocellulosic content, while the food waste used in this study had high concentrations of proteins and lipids (Table S1). Therefore, the relative abundance and activity of genera involved in amino acid and lipid metabolism (i.e. *Acidaminococcus* and *Lachnospiraceae NK3A20*) was higher in the rumen AnDMBR compared to in the rumen inoculum.^70,82^ Some populations in the rumen AnDMBR, like *Fibrobacter*, *Megasphaera*, and *Methanobrevibacter*, were more active in the dynamic membrane than in the bulk suspended biomass (Figures S5, S10, and S12), consistent with a few studies focusing on methanogenesis or chain elongation^46,77,83^. However, there was no significant difference in the microbial community structure and activity of the bulk suspended biomass and dynamic membrane, likely because the dynamic membrane was disrupted every hour and mixed with the bulk solution (Figures 2 and 4).

**Figure 2.**
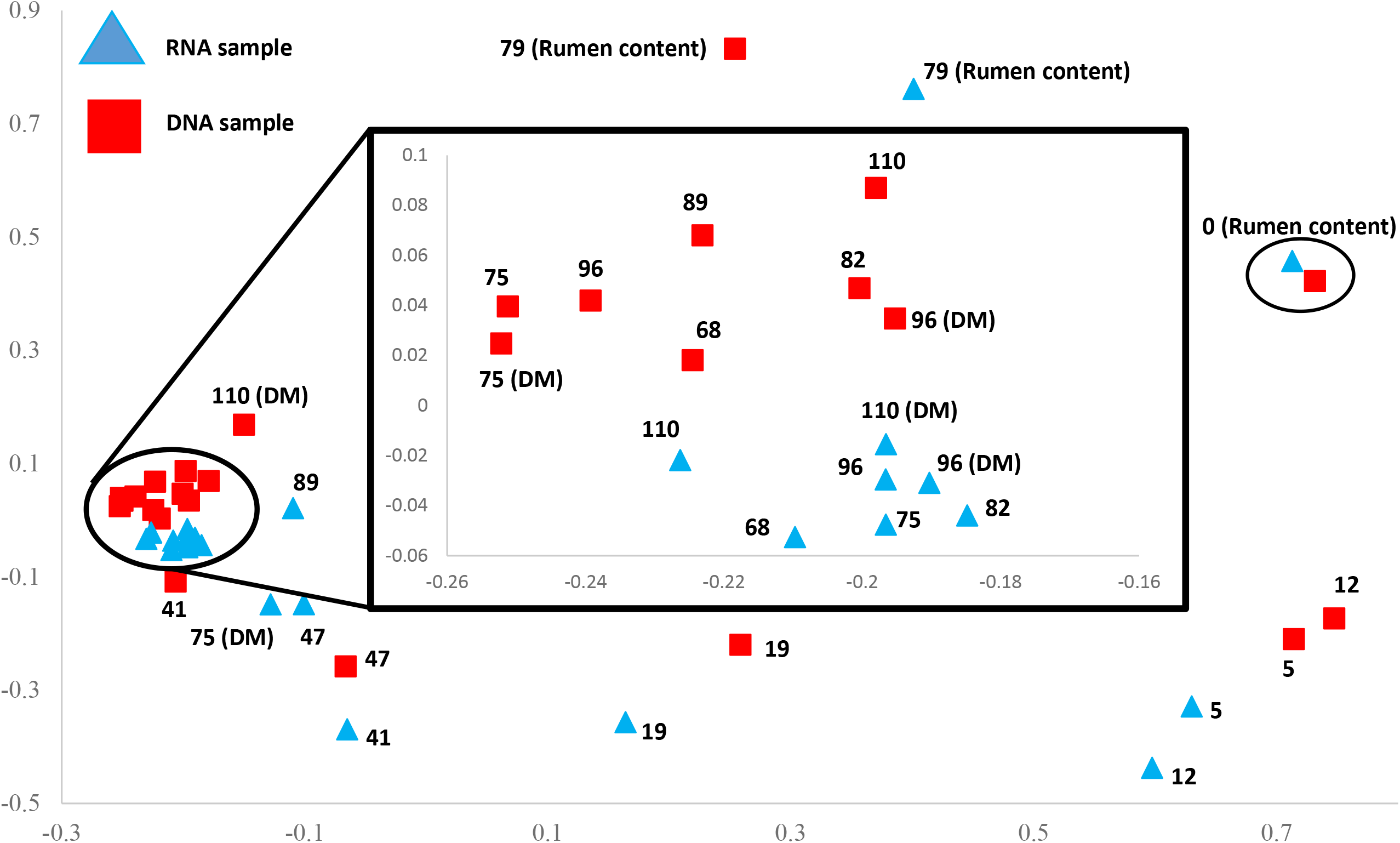
Non-metric multidimensional scaling (NMDS) plot based on Bray-Curtis distance of microbial community present (16S rRNA gene) and active (16S rRNA) in rumen inoculum and suspended biomass samples over time. The numbers represent the days of sampling. DM indicates that the samples were taken from the dynamic membrane.

This is the first study to enhance the hydrolysis and VFA production from food waste using a rumen-based AnDMBR and to provide insight in the microbial community present and active in the bioreactor over time utilizing DNA and RNA sequencing. Although the substrate and engineering considerations deviates the bioreactor from 100% replicating a cow rumen, recreating the rumen HRT, SRT, temperature and regurgitation achieved high NDF, ADF removal rates and VFA yields. Up to 66.7% of active microbial community in the bioreactor can be traced back to the rumen, including important hydrolytic bacteria like *Fibrobacter*, *R. flavefaciens* and *R. albus*. The microbial community in the rumen reactor is resilient enough to quickly recover their composition and performance after an operational disruption and re-inoculation. The conditions used here allowed us to developed a new type of membrane bioreactor that is easy to operate (the system did not present problems related to fouling), able to produce VFA faster than other systems (low reactor footprint), resilient and versatile.

## SUPPORTING INFORMATION

## ACKNOWLEDGEMENTS

The authors would like to thank Steve Donajkowski and Ethan Kennedy for their assistance with the bioreactor system. This work was supported by REFRESCH (Global Challenges for the Third Century program, Office of the Provost, University of Michigan) and the U.S. National Science Foundation (Sustainability Research Networks 1444745). Shilva Shrestha was supported by an Integrated Training in Microbial Systems Fellowship funded by the Boroughs Wellcome Fund, the University of Michigan Rackham Predoctoral Fellowship, and a Water Environment Federation Canham Graduate Studies Scholarship.

