## Supporting information for "Simulating Rumen Conditions using an Anaerobic Dynamic Membrane Bioreactor to Enhance Hydrolysis of Lignocellulosic Biomass"

The supporting information has 21 pages including 13 figures and 5 tables.

**Table S1.** Characteristics of inoculum and substrate

| Parameter | Inoculum (rumen content) | Substrate (food waste) | Parameter | Inoculum (rumen content) | Substrate (food waste) |
|---|---|---|---|---|---|
| Total solids (TS) (g $L^{-1}$) | 58.9 ± 1.6 | 72.6 ± 4.3 | Volatile fatty acids (VFA) (g VFA as COD $L^{-1}$) | 16.1 ± 0.4 | 0.9 ± 0.2 |
| Volatile solids (VS) (g $L^{-1}$) | 48.3 ± 1.5 | 68.9 ± 5.2 | Acetate (g COD $L^{-1}$) | 7.4 ± 0.1 | 0.9 ± 0.2 |
| Total suspended solids (TSS) (g $L^{-1}$) | 46.5 ± 0.9 | 49.2 ± 8.4 | Propionate (g COD $L^{-1}$) | 5.1 ± 0.4 | ND |
| Volatile suspended solids (VSS) (g $L^{-1}$) | 39.6 ± 0.5 | 48.0 ± 7.9 | Isobutyrate (g COD $L^{-1}$) | ND | ND |
| pH | 5.7 ± 0.3 | 4.5 ± 0.3 | Butyrate (g COD $L^{-1}$) | 3.6 ± 0.1 | ND |
| Partial alkalinity (g $CaCO_3$ $L^{-1}$) | 0.9 ± 0.5 | NA | Isovalerate (g COD $L^{-1}$) | ND | ND |
| Total alkalinity (g $CaCO_3$ $L^{-1}$) | 5.4 ± 0.8 | 0.2 ± 0.2 | Valerate (g COD $L^{-1}$) | ND | ND |

| | | | | | |
|---|---|---|---|---|---|
| Soluble chemical oxygen demand (sCOD) (g COD $L^{-1}$) | 22.7 ± 0.7 | 30.4 ± 5.8 | Caproate (g COD $L^{-1}$) | ND | ND |
| Proteins (g $L^{-1}$) | NA | 23.9 ± 0.4 | Heptanoate (g COD $L^{-1}$) | ND | ND |
| Lipids (g $L^{-1}$) | NA | 7.5 ± 0.1 | Octanoate (g COD $L^{-1}$) | ND | ND |
| Neutral detergent fiber (NDF) (g $L^{-1}$) | 7.3 ± 0.7 | 11.3 ± 1.5 | Acid detergent fiber (ADF) (g $L^{-1}$) | 3.7 ± 0.5 | 9.0 ± 1.6 |

All data are reported as mean ± standard deviation. The mean and standard deviation for the rumen content were calculated using three technical replicates of the samples collected on day 0 and day 79. The data for the inoculum are for the mixture of solid and liquid rumen content fractions used to inoculate the system on day 0 and day 79. The data for the substrate are for the diluted food waste fed to the rumen AnDMBR. The mean and standard deviation for food waste were calculated using three technical replicates analyzed for all the batches used.

ND: Not detected, NA: Not applicable

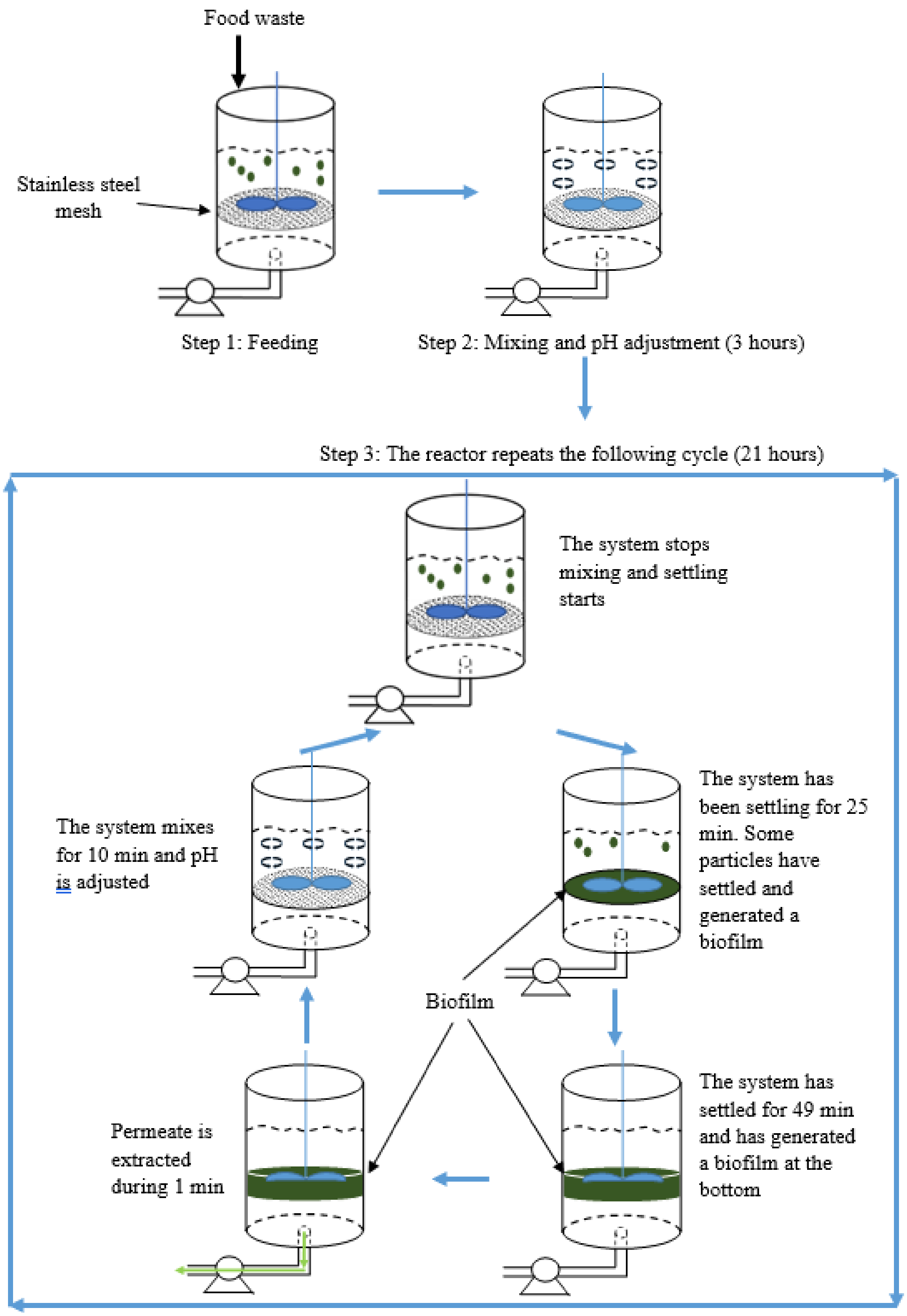


**Figure S1.** Schematic representation of the rumen AnDMBr

**Table S2.** Rumen reactor performance

| Parameter | Rumen reactor | Parameter | Rumen reactor |
|---|---|---|---|
| VFA yield (g VFA as COD g $VS_{fed}^{-1}$) | 0.55 ± 0.12 | *VFA (g VFA as COD $L^{-1}$) | 6.0 ± 1.9 |
| Methane yield (gCH4 as COD $gCOD_{fed}^{-1}$) | 2.5 ± 1.2 | *Acetate (g COD $L^{-1}$) | 2.1 ± 0.9 |
| $CH_4$ content in biogas (%) | 53.1 ± 9.7 | *Propionate (g COD $L^{-1}$) | 1.9 ± 0.7 |
| $CO_2$ content in biogas (%) | 39.7 ± 9.7 | *Isobutyrate (g COD $L^{-1}$) | ND |
| $H_2$ content in biogas (%) | 6.8 ± 12.2 | *Butyrate (g COD $L^{-1}$) | 1.1 ± 0.6 |
| *g VFA as COD g $sCOD^{-1}$ | 0.84 ± 0.12 | *Isovalerate (g COD $L^{-1}$) | 0.1 ± 0.2 |
| Partial alkalinity (g $CaCO_3$ $L^{-1}$) | 0.31 ± 0.25 | *Valerate (g COD $L^{-1}$) | 0.6 ± 0.3 |
| Total alkalinity (g $CaCO_3$ $L^{-1}$) | 1.57 ± 0.67 | *Caproate (g COD $L^{-1}$) | 0.1 ± 0.2 |
| NDF removal (%) | 58.9 ± 8.4 | *Heptanoate (g COD $L^{-1}$) | ND |
| ADF removal (%) | 69.0 ± 8.6 | *Octanoate (g COD $L^{-1}$) | ND |
| §Protein removal (%) | 42.0 ± 10.8 | §Lipid removal (%) | 90.0 ± 3.8 |

All data are reported as mean ± standard deviation for the 110 days of operation. *Concentrations measured in the permeate. § Proteins and lipids in the reactor permeate were only measured on days 90, 95, 100, 105, and 107.

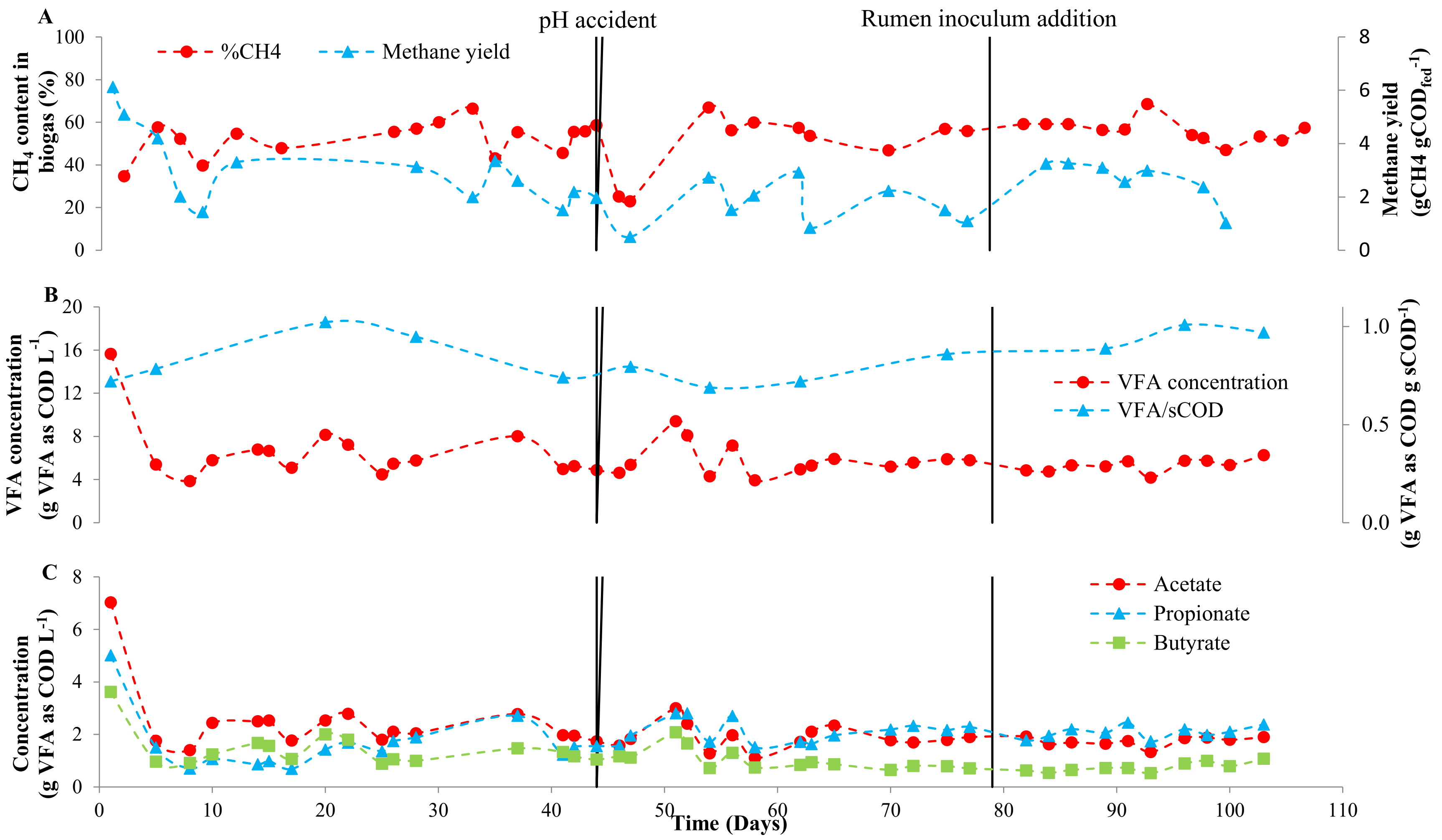


**Figure S2.** Methane yield and methane content in biogas (A) and VFA concentration and VFA over sCOD ratio (B) and acetate, propionate, and butyrate concentrations (C). The vertical line indicates the increase of pH for four hours (day 44) and the addition of rumen inoculum (day 79).

**Table S3**. Performance of other rumen-based bioreactors (part A)

| Studies | Substrate | NDF removal (%) | ADF removal (%) | Cellulose removal (%) | Hemicellulose removal (%) | VFA yield |
|---|---|---|---|---|---|---|
| This study | Food waste | 58.9 | 69.0 | ND | ND | 0.55 ± 0.1 g VFA as COD g $VS_{fed}^{-1}$<br>0.4 ± 0.1 g VFA g $VS_{fed}^{-1}$<br>5.7 ± 0.6 mmol VFA $gVS_{fed}^{-1}$ |
| 1 | Maize silage and concentrate | ND | ND | ND | ND | §0.22 g VFA g $VS_{fed}^{-1}$ |
| 2 | Corn stover and food waste | ND | ND | 65 | 43 | 0.4 g VFA as COD g $VS_{fed}^{-1}$ |
| 3 | Corn stover | ND | ND | 31 | 31 | §0.22 ± 0.1 g VFA g $VS_{fed}^{-1}$ |
| 4 | Cellulose | ND | ND | ND | ND | 0.32 g VFA as COD g $VS_{fed}^{-1}$ |
| 5* | Corn stover | 66.8 | ND | ND | ND | §0.4 ± 0.1 g VFA g $VS_{fed}^{-1}$ |

| | | | | | | |
|---|---|---|---|---|---|---|
| 6* | Municipal solid wastes and alfalfa | 76 | 73 | ND | ND | §3.2 mmol VFA g $VS_{fed}^{-1}$ |
| 7* | Barley straw, rye straw, and Maize stover | 57 | ND | 55 | 59 | §2.2 mmol VFA g $VS_{fed}^{-1}$ |

ND: Non determined. *These studies used the same reactor configuration §The values provided here were calculated using information from the paper

**Table S3**. Performance of other rumen-based bioreactors (part B)

| Studies | SRT (days) | HRT (days) | OLR (g VS $L_R^{-1}$ $day^{-1}$) | Mesh pore size (µm) |
|---|---|---|---|---|
| This study | 4.0 ± 1.6 (TSS based)<br>2.1 ± 0.6 (TS based) | 0.50 ± 0.1 | 18 ± 2 | 100 |
| 1 | 20 | 2.375 | 1.14 (g COD $L_R^{-1}$ $day^{-1}$) | 0.04 |
| 2 | unknown | 33 | 6.2 (g COD $L_R^{-1}$ $day^{-1}$) | 50 |
| 3 | 31 | 31 | | No membrane |
| 4 | 7 | 0.67 | 22 | No membrane |
| 5* | 4 | 0.75 | 10 | 30 |
| 6* | 3.8 | 0.5 | 19.3 | 30 |
| 7* | 3 | 0.5 | 17.7 | 30 |

ND: Non determined. *These studies used the same reactor configuration

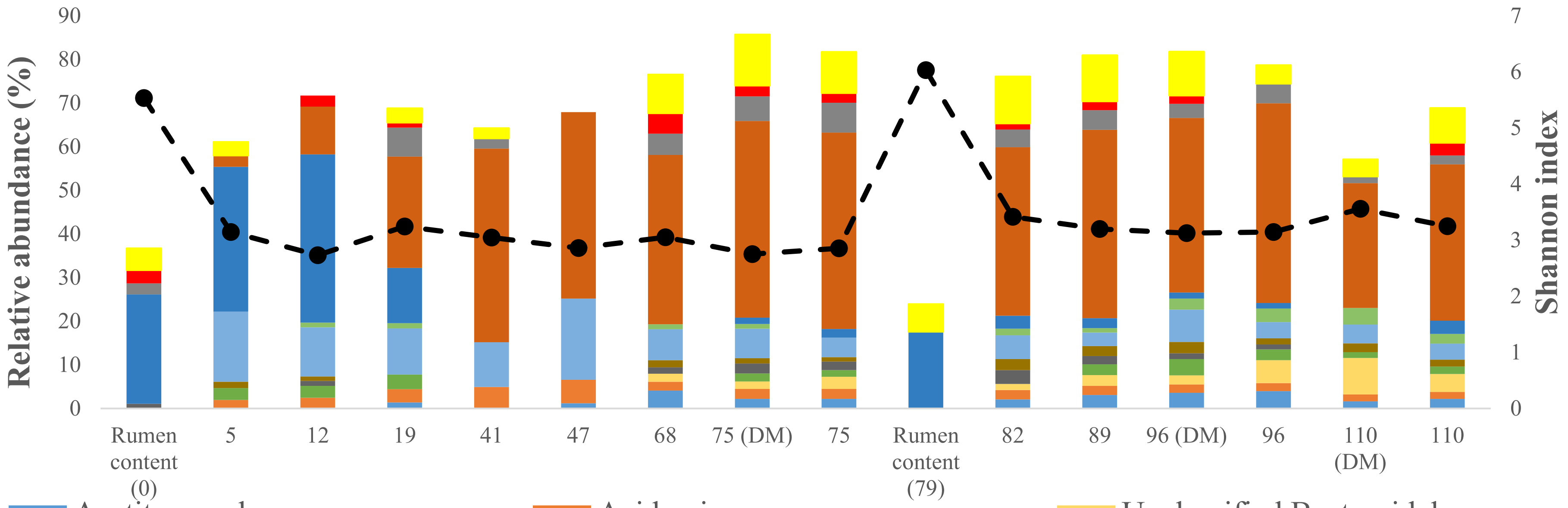


**Figure S3**. Relative abundance (determined by 16S rRNA gene sequencing) of dominant genera in the inoculum and bioreactor suspended biomass over time. Only genera present at relative activities greater than 1% in at least 60% of the samples (n=10) or present at a relative activity of 15% at least once are shown. The Shannon diversity index was calculated considering all the ASVs. DM indicates that the samples were taken from the dynamic membrane.

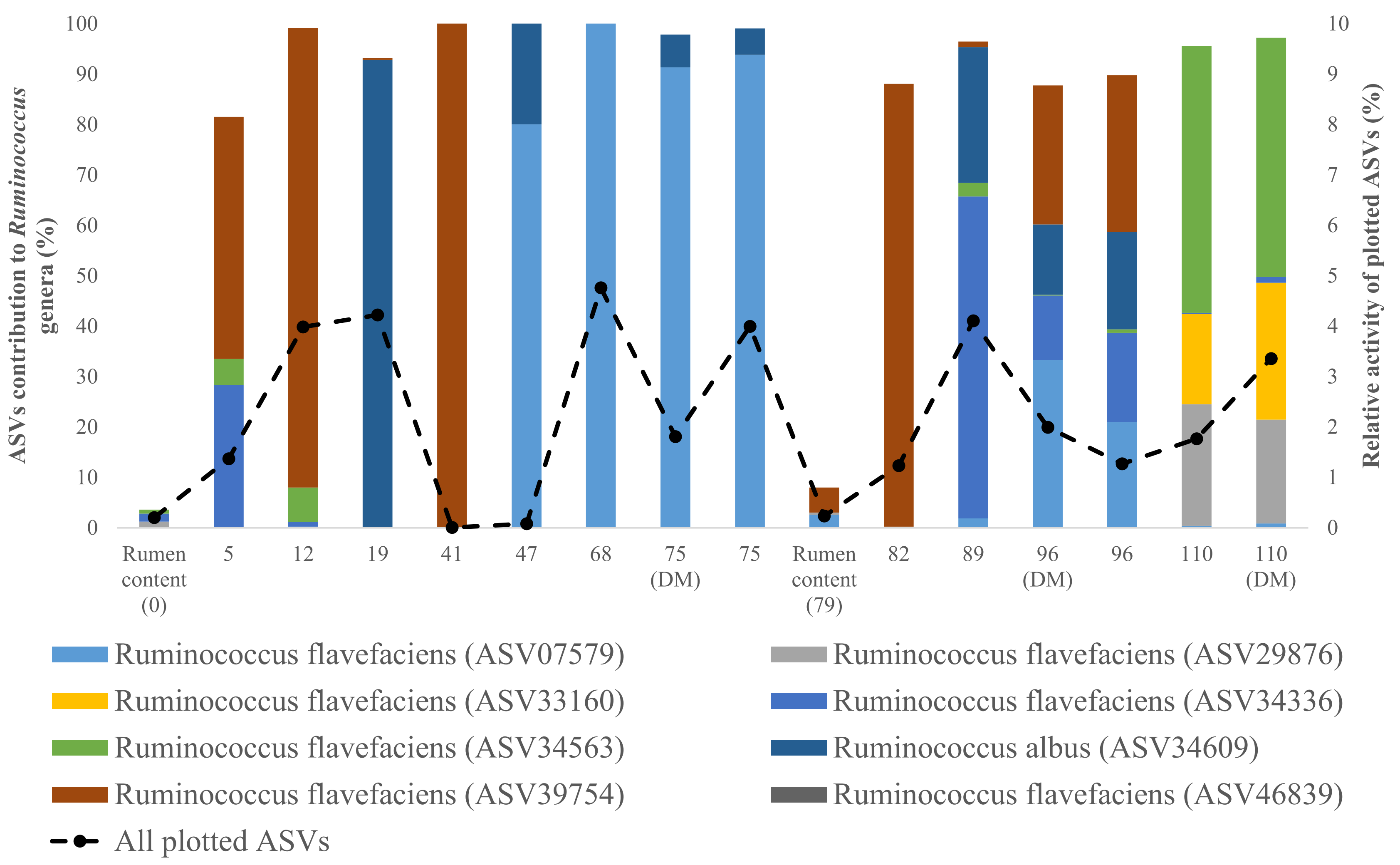


**Figure S4.** Relative activity of ASVs making up the *Ruminococcus* genera and relative activity the sum of all the ASV plotted. DM indicates that the samples were taken from the dynamic membrane.

**Table S4.** Species assignation to ASVs making up *Ruminococcus* genera

| BLAST values | Max score | Total score | Query cover (%) | E value | Percent identity (%) | Accession length |
|---|---|---|---|---|---|---|
| *R. flavefaciens* (ASV 07579) | 468 | 468 | 100 | 2E-127 | 100 | 1519 |
| *R. flavefaciens* (ASV 29876) | 429 | 429 | 100 | 1E-115 | 97 | 728 |
| *R. flavefaciens* (ASV 33160) | 457 | 457 | 100 | 5E-124 | 99 | 253 |
| *R. flavefaciens* (ASV 34336) | 468 | 464 | 100 | 2E-127 | 100 | 1505 |
| *R. flavefaciens* (ASV 34563) | 440 | 440 | 100 | 5E-119 | 98 | 728 |
| *R. albus* (ASV 34609) | 468 | 468 | 100 | 2E-127 | 100 | 1529 |
| *R. flavefaciens* (ASV 39754) | 468 | 468 | 100 | 2E-127 | 100 | 1529 |
| *R. flavefaciens* (ASV 46839) | 462 | 462 | 100 | 1E-125 | 99 | 1464 |

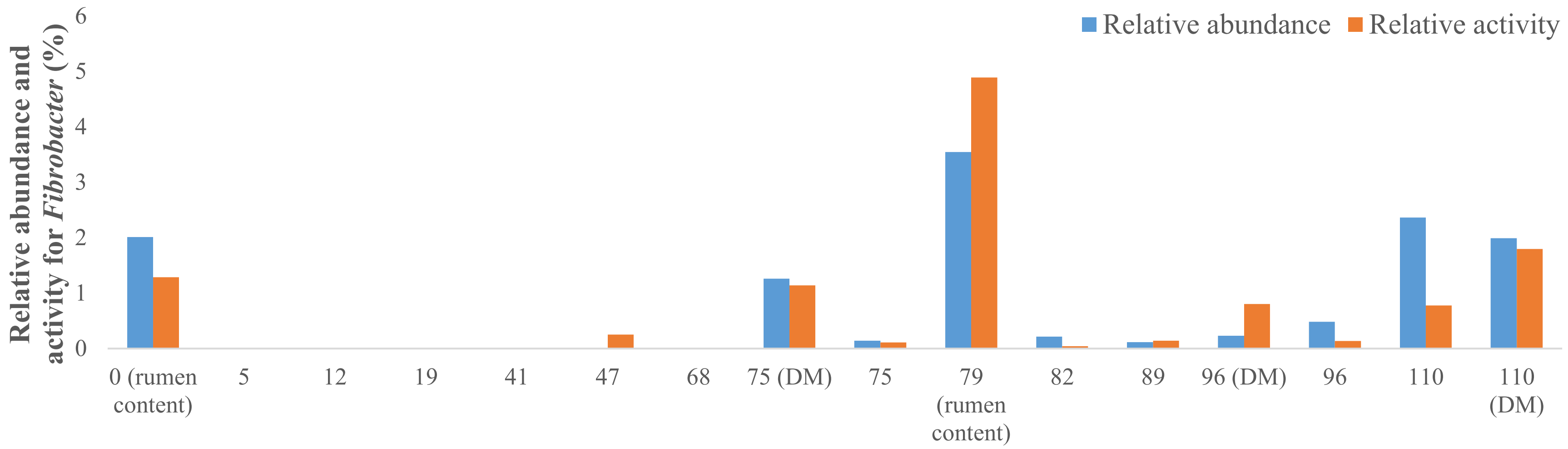


**Figure S5.** Relative activity and abundance of *Fibrobacter* in the rumen AnDMBr. DM indicates that the samples were taken from the dynamic membrane.

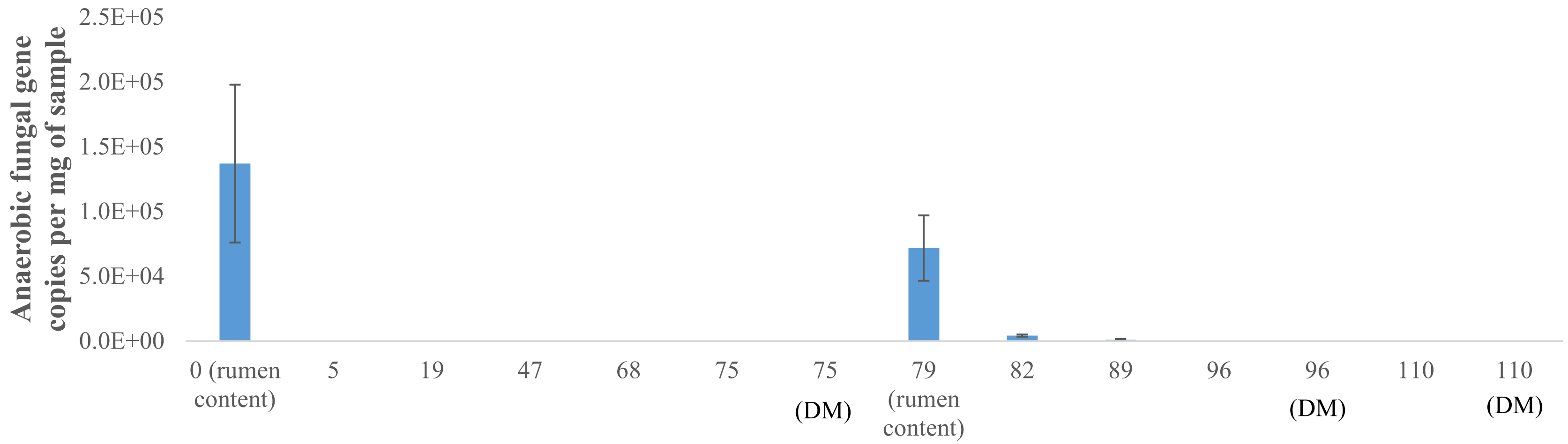


**Figure S4.** Number of anaerobic fungal SSU rRNA gene copies detected per mg of sample. Error bars indicating standard deviation.

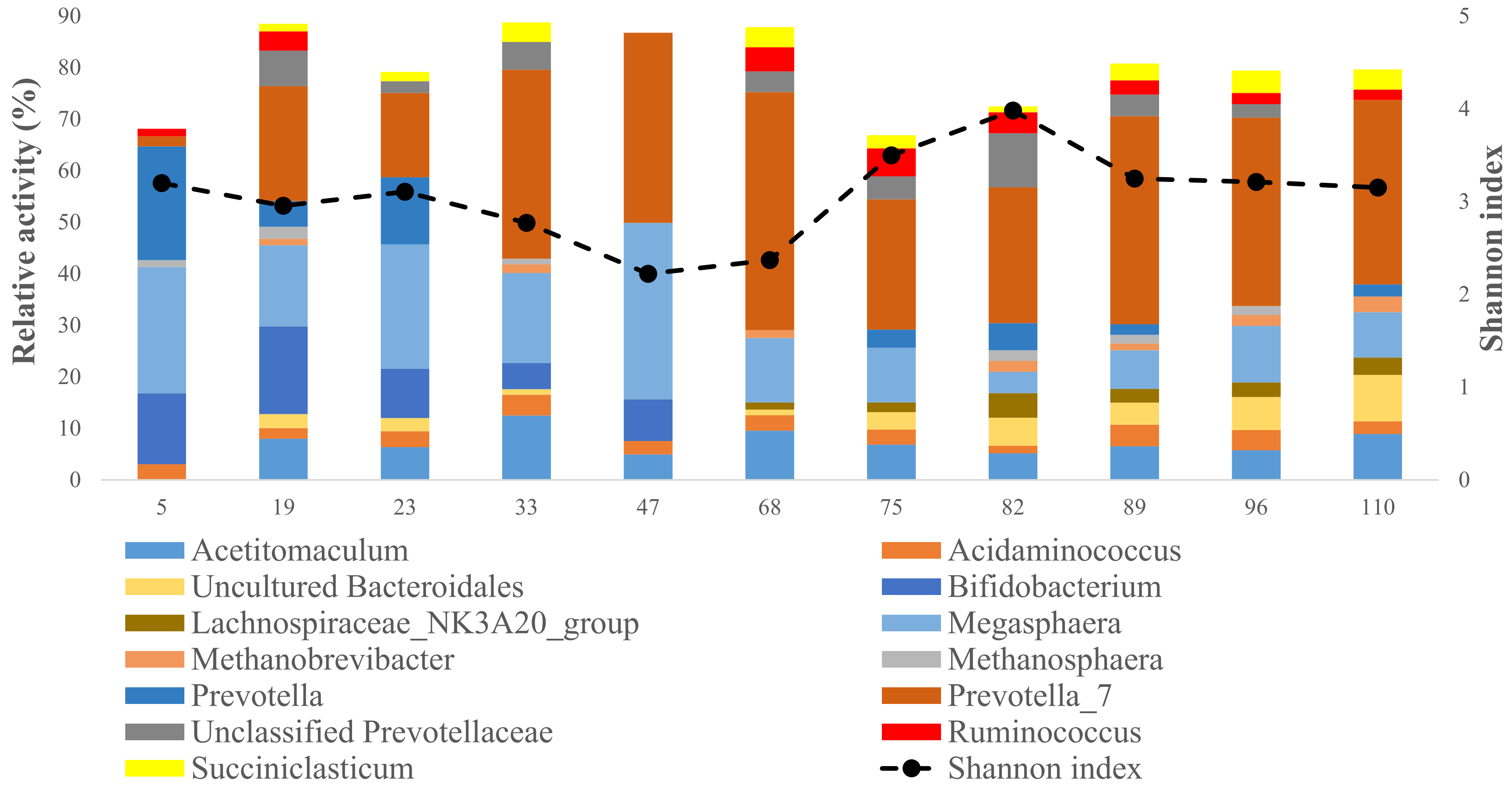


**Figure S7.** Relative activities (determined by 16S rRNA sequencing) of dominant genera in permeate over time. Only genera present at relative activities greater than 1% in at least 60% of the samples or present at a relative activity of 15% at least once are shown. The Shannon diversity index was calculated considering all the ASVs. DM indicates that the samples were taken from the dynamic membrane.

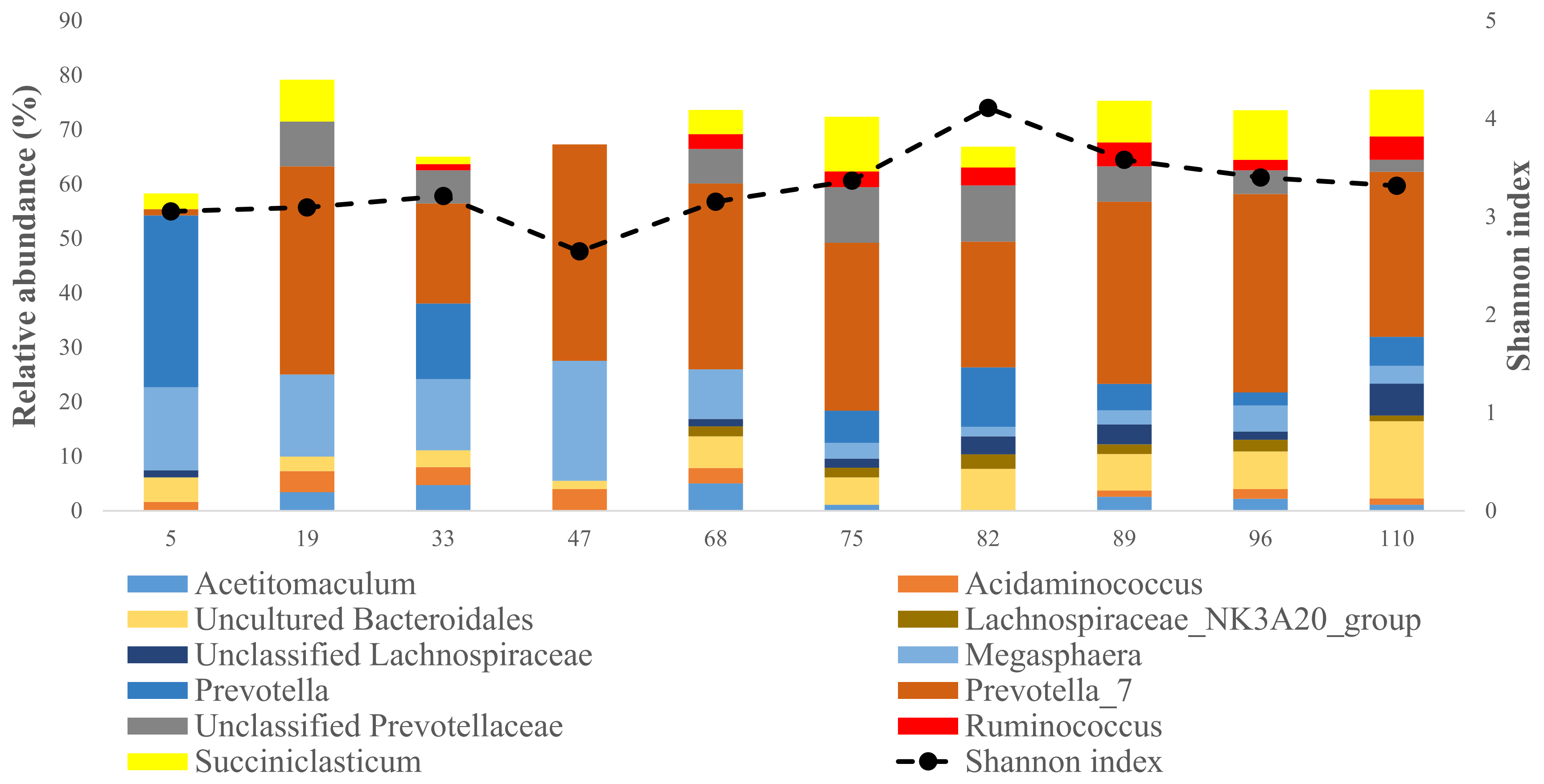


**Figure S8.** Relative abundance (determined by 16S rRNA gene sequencing) of dominant genera in permeate over time. Only genera present at relative activities greater than 1% in at least 60% of the samples or present at a relative activity of 15% at least once are shown. The Shannon diversity index was calculated considering all the ASVs. DM indicates that the samples were taken from the dynamic membrane.

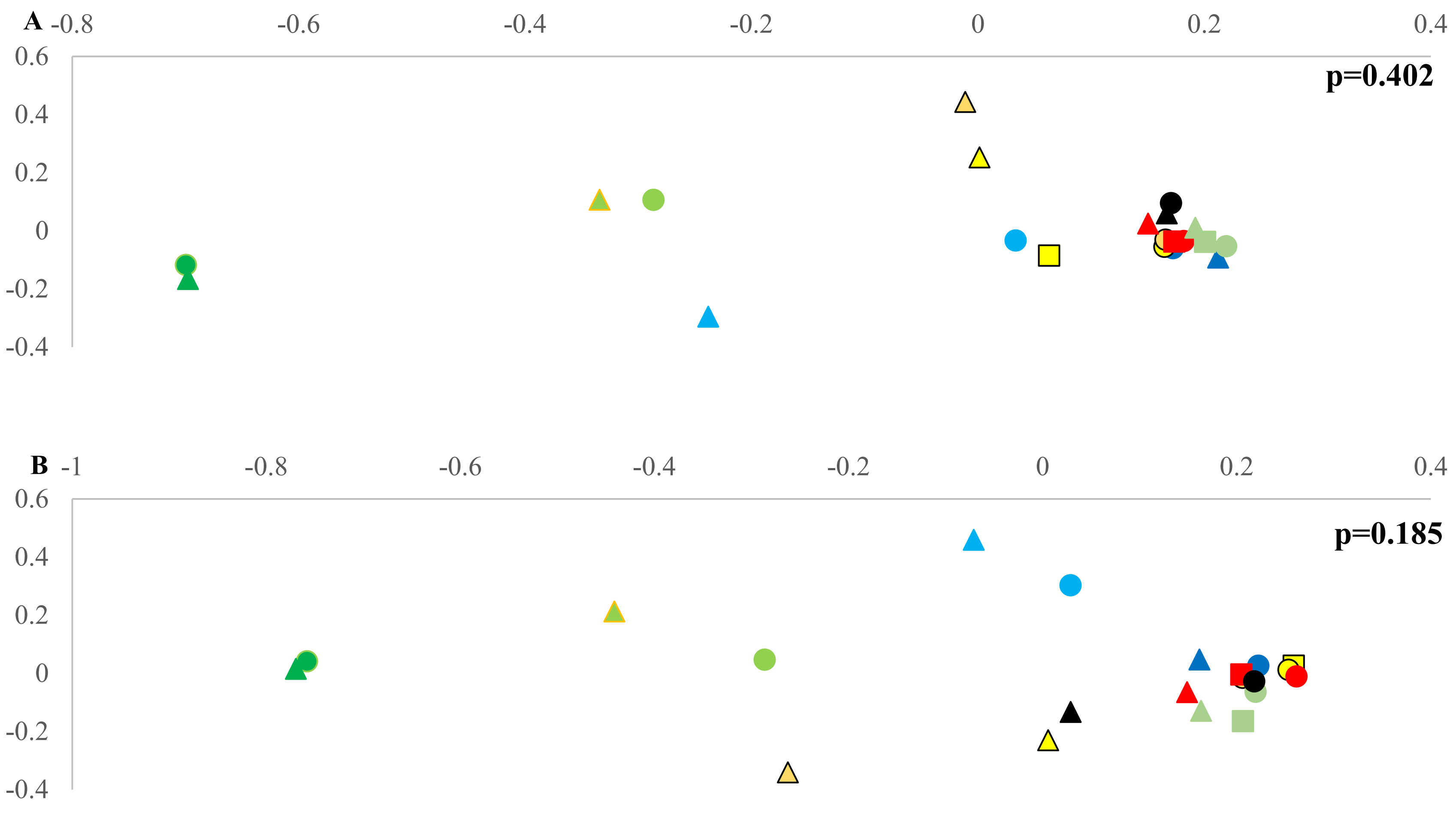


**Figure S9.** Comparison of microbial community dissimilarities based on analysis of molecular variance for 16S rRNA (A) and 16S rRNA gene. The triangles represent permeate samples, the circles represent suspended biomass samples and the squares represent samples collected from the dynamic membrane. The samples collected on the same have the same colors.

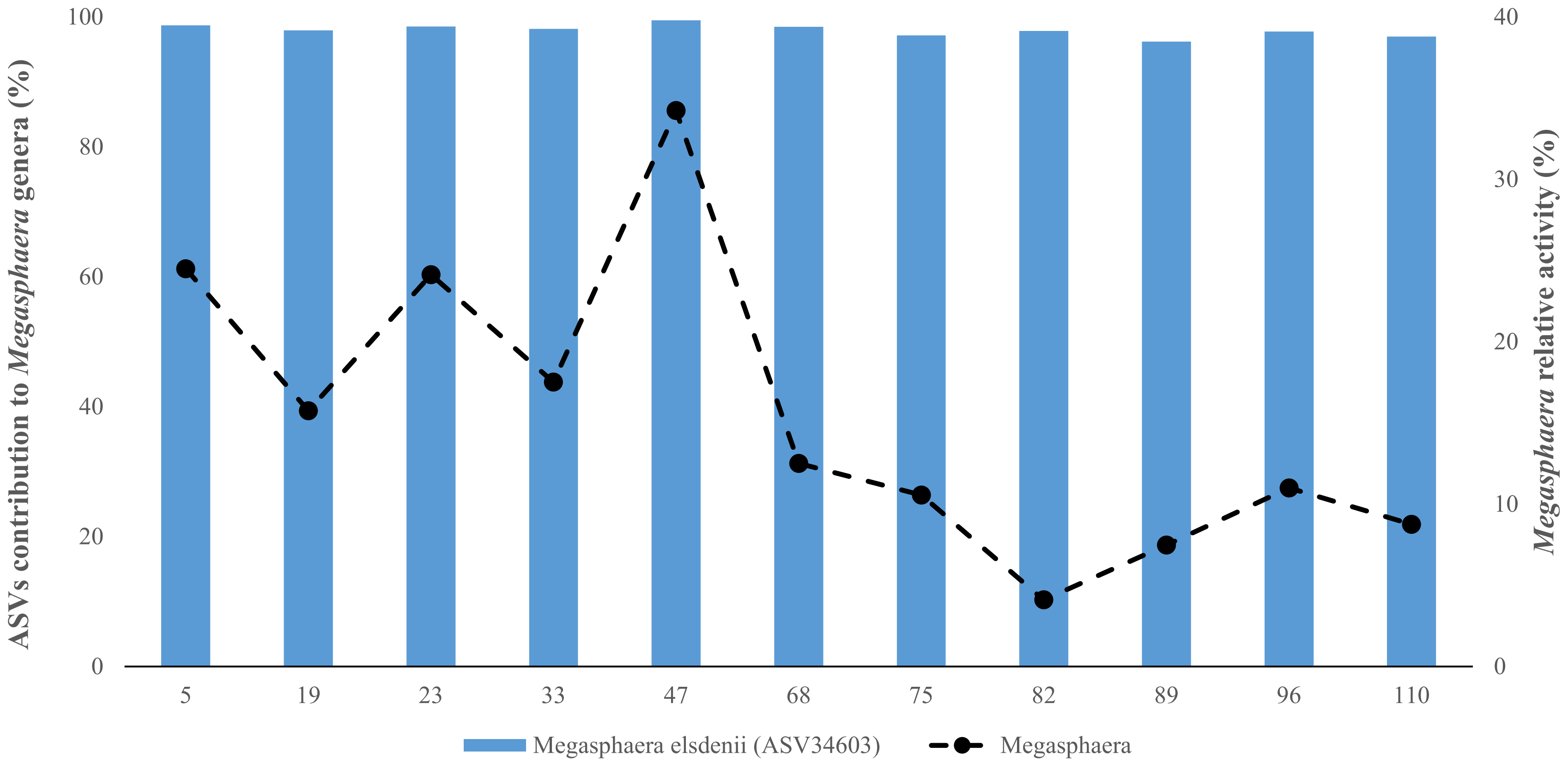


**Figure S10.** Relative activity of ASVs making up the *Megasphaera* genus and relative activity of the genus *Megasphaera* in the permeate samples

**Table S5**: Species assignation to ASVs making up *Megasphaera* genera

| BLAST values | Max score | Total score | Query cover (%) | E value | Percent identity (%) | Accession length |
|---|---|---|---|---|---|---|
| *Megasphaera elsdenii* (ASV34603) | 466 | 466 | 100 | 8E-127 | 100 | 1458 |

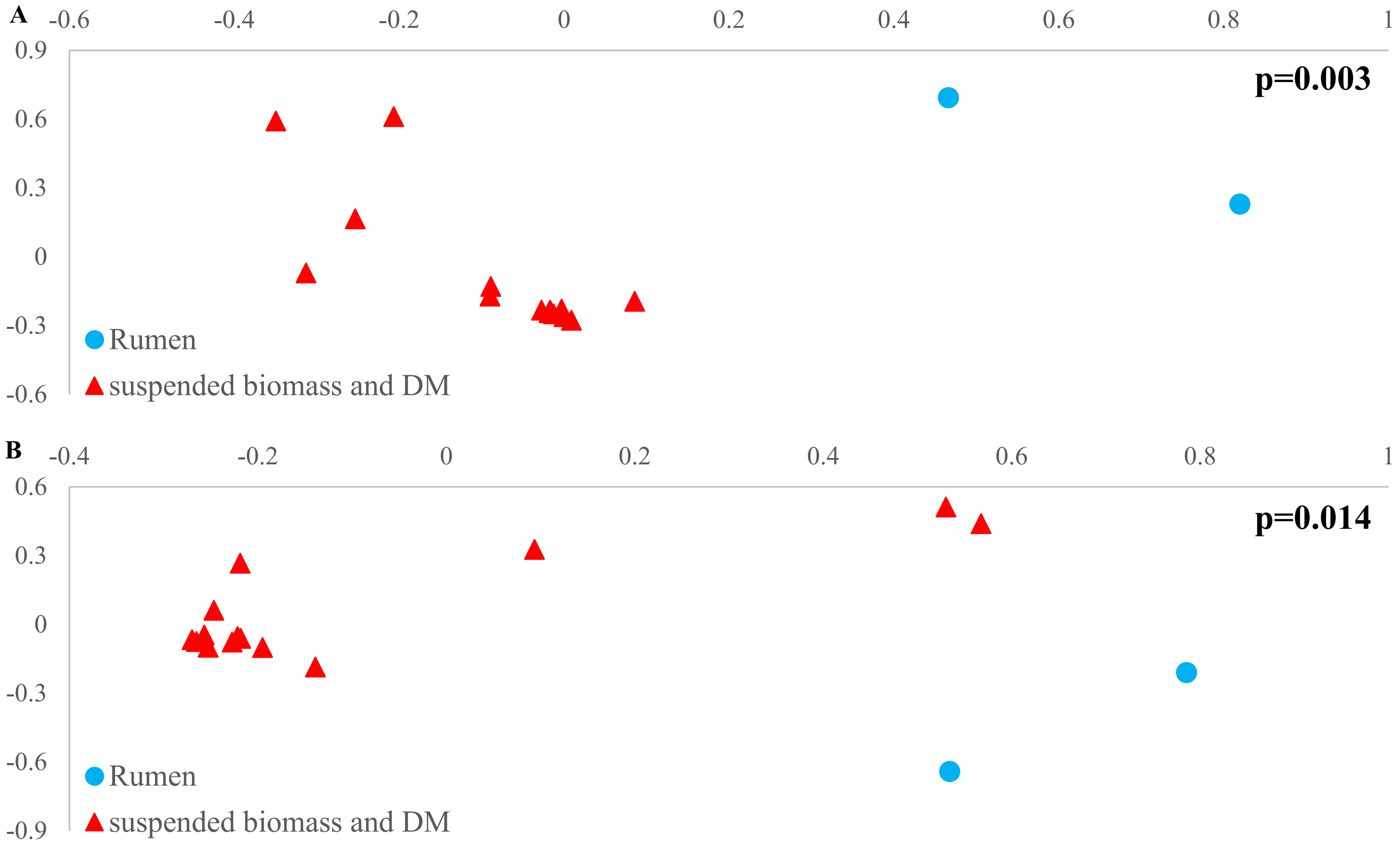


**Figure S11.** Comparison of microbial community dissimilarities based on analysis of molecular variance for 16S rRNA (A) and 16S rRNA gene (B). DM stands for dynamic membrane.

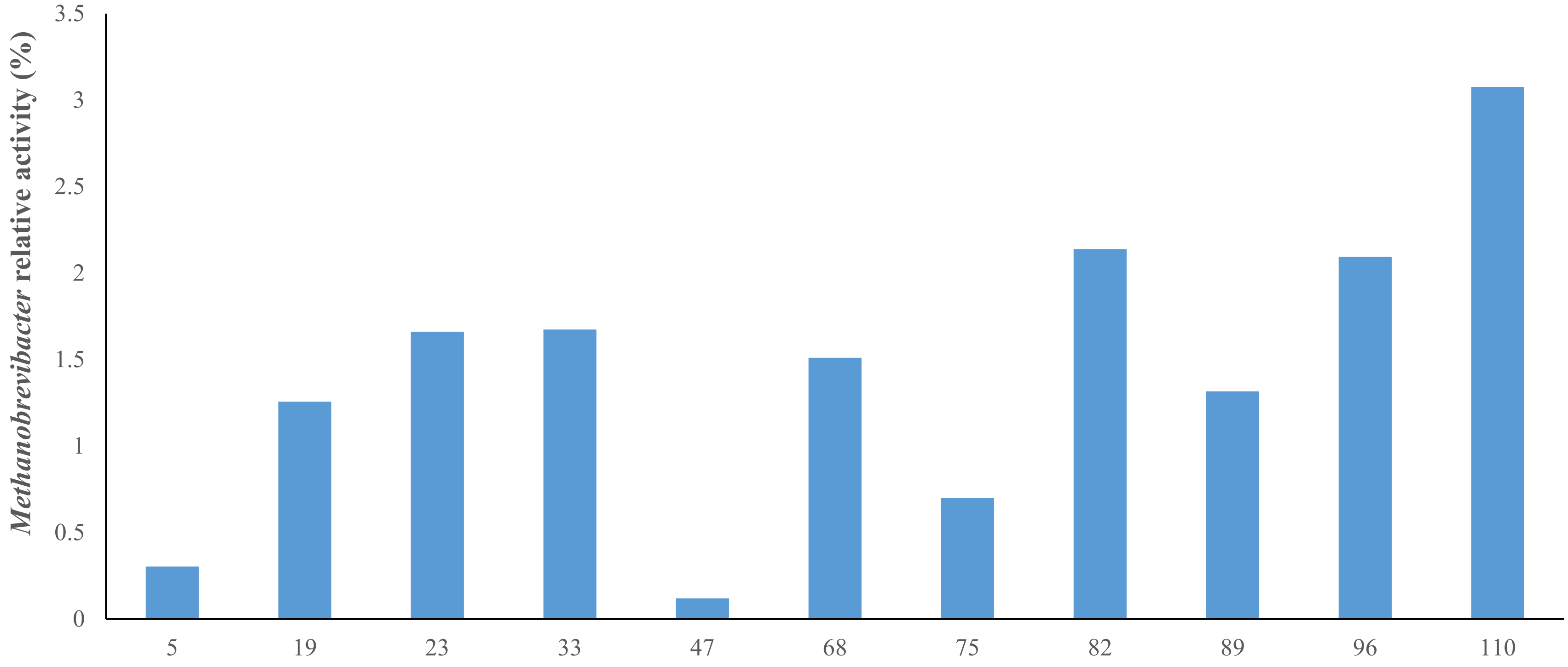


**Figure S12.** Relative activity of *Methanobrevibacter* in the rumen AnDMBR permeate.

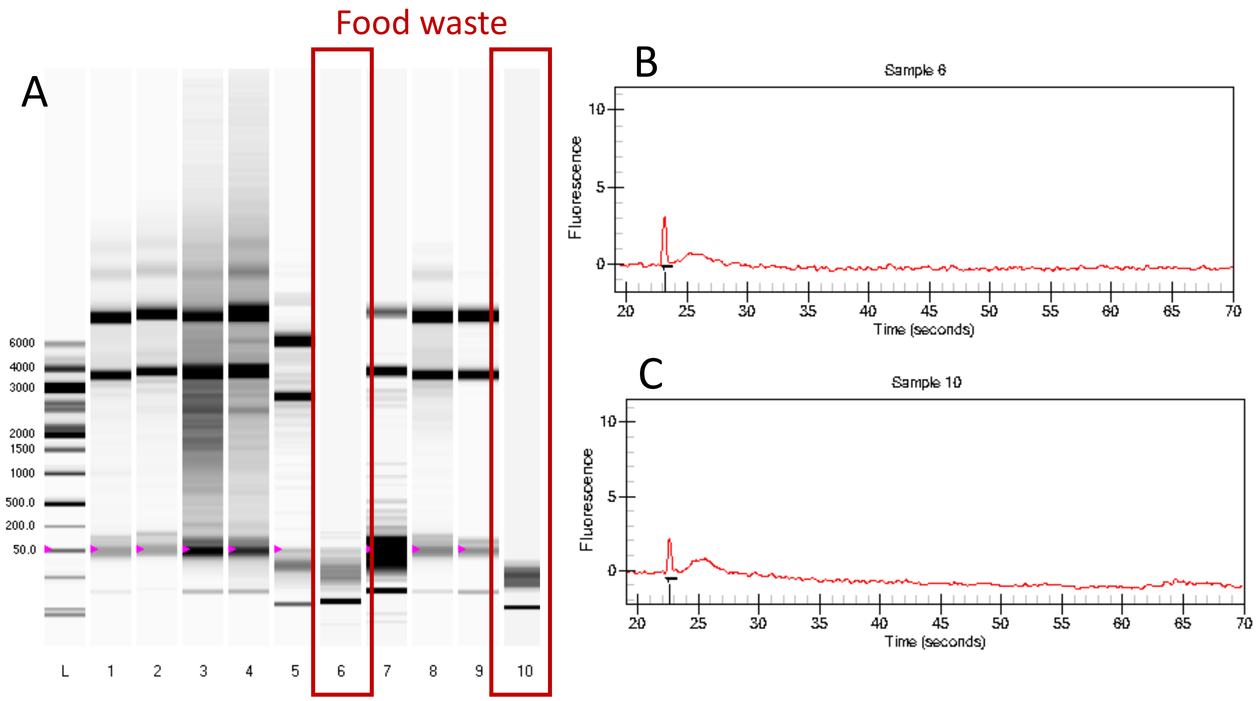


**Figure S13.** Electrophoresis image of RNA extracted from the AnDMBR permeate (Lanes 1, 2, and 5), AnDMBR bulk solution (Lanes 4, 8, and 9), dynamic membrane (Lane 7), rumen inoculum (Lane 3), and food waste (Lanes 6 and 10) (A) and fluorescence intensities (B and C) of food waste corresponding to lanes 6 and 10 (A) taken with a Bioanalyzer.
